# Soil fungi remain active and invest in storage compounds during drought independent of future climate conditions

**DOI:** 10.1101/2023.10.23.563577

**Authors:** Alberto Canarini, Lucia Fuchslueger, Jörg Schnecker, Dennis Metze, Daniel B. Nelson, Ansgar Kahmen, Margarete Watzka, Erich M. Pötsch, Andreas Schaumberger, Michael Bahn, Andreas Richter

## Abstract

Microbial growth is central to soil carbon cycling. However, how microbial communities grow under climate change is still largely unexplored. In an experiment simulating future climate conditions (increased atmospheric CO_2_ and temperature) and drought, we traced ^2^H or ^18^O applied via water-vapor exchange into fatty acids or DNA, respectively, allowing to measure community- and group-level adjustments in soil microbial physiology (replication, storage product synthesis, and carbon use efficiency, CUE). We show, that while overall community-level growth decreased by half during drought, fungal growth remained stable demonstrating an astonishing resistance of fungal activity against soil moisture changes. In addition, fungal investment into storage triglycerides increased more than five-fold under drought. CUE (the balance between anabolism and catabolism) was unaffected by drought but decreased in future climate conditions. Our results highlight that accounting for different growth strategies can foster our understanding of soil microbial contribution to C cycling and feedback to climate change.

## 1. Introduction

Plant inputs and soil organic matter decomposition have been considered the main drivers controlling the soil organic carbon (SOC) balance, but more recently, growth and biomass production rates of soil microbial communities have been identified as main contributors to SOC formation and persistence (*1–3*). In addition, the efficiency at which microbes allocate C to growth relative to respiratory processes (i.e., microbial carbon use efficiency or CUE) has been suggested to be a key parameter for predicting SOC stock and can strengthen prediction of SOC feedback in response to a changing climate (*4*). Improving our ability to accurately quantify soil microbial growth is essential to understand the controls on soil carbon (C) and nutrient cycling (*5*, *6*) and generally the mechanisms behind terrestrial carbon cycling.

Because soil microbial communities respond sensitively to climate change, shifts in microbial physiology can cause large repercussions on global C and nutrient cycles (*6–8*). Elevated CO_2_ (eCO_2_) induces mostly indirect effects on soil microbes by stimulating primary productivity (*9*) and thus increasing C resources available for heterotrophic soil microbes, leading to enhanced microbial growth and decomposition rates (*10–12*). Warming can increase plant productivity, but also directly stimulate microbial physiological activity, decreasing microbial CUE and accelerating soil carbon losses (*6*, *7*, *13*). Drought, in contrast, reduces accessibility to C and nutrients, and consequently can decrease soil microbial activity (*14*, *15*) to the point, where in the absence of water, microbes become dormant or die. On an ecosystem level, drought can lead to soil C losses (*16*), as soil respiration is generally less sensitive to dry conditions compared to plant primary productivity (*17*, *18*). Furthermore, the impact of drought on soil respiration and primary productivity can be modified by eCO_2_ and warming (*19–21*). Assessing microbial growth responses to drought, and possible interactions with other global change factors in field experiments is therefore crucial. Yet, multifactorial climate change experiments conducted under field conditions measuring soil functions and microbial communities are rare (*22*), rendering interactive effects on soil microbial feedbacks largely unexplored.

It is notoriously challenging to quantify soil microbial growth, and even more so under drought conditions, without the application of substrate or water (*23*). Over the last decade, advances in substrate-free, ^18^O-based stable isotope probing (^18^O-SIP), tracing ^18^O assimilation into newly produced DNA, improved quantitative measures of physiological rates of microbes on a community-level (*24*), as well as of taxon-specific bacterial growth rates (*25*, *26*). Recently it was shown that it is possible to enrich soil water with ^18^O indirectly via water vapor equilibration, avoiding direct water additions to the soil. This now allows the quantification of soil microbial community-level and taxon-level growth rates of bacteria during drought conditions (*27*, *28*). The link between growth rates and microbial identity is a major focus in microbial ecology and can provide mechanistic insights into the role of soil microbial taxa and community dynamics for biogeochemical C and nutrient cycling. Microbial biomass can increase as a result of cell replication, but also through the synthesis of storage compounds (*29*). The ability of microbial populations to invest resources in storage enables them to detach their metabolic activity from the immediate resource availability, thereby facilitating a wider range of microbial responses to environmental fluctuations (*30*). Quantifying taxon-specific growth rates via DNA (or RNA) based method (^18^O-SIP) has been mainly applied to link bacterial community structure to activity rates. However, it requires a separate analysis to measure fungal activity, which has been largely overlooked. At the same time ^18^O-SIP yields a high number of samples to process and analyze (*31*, *32*) which makes it challenging to apply to large sample numbers.

Deuterium (^2^H) incorporation into fatty acids can serve as an alternative SIP approach to measure microbial growth, as all microorganisms synthesize lipids and fatty acids to build and maintain cell membranes regardless of their metabolic activity and cell cycle stage. ^2^H-labelling has been used in single cell studies (*33*), pure cultures (*34*) and soil microbial communities (*35*). Lipids likely require less investment into repair mechanisms compared to nucleic acids or proteins, which can be resynthesized during repair and cellular maintenance potentially affecting growth rates calculation (*35*, *36*), and provide a precise measure of microbial membrane production rates and cell growth. The major advantages of tracing ^2^H into fatty acids (^2^H-PLFA-SIP) are that it allows the simultaneous and sensitive quantification of bacterial and fungal replication rates (*37*) and the concurrent measurement of storage compound production (*38*). Although compared to DNA/RNA-based methods, PLFA analysis has a lower taxonomic resolution, it provides a more robust and quantitative estimate of microbial biomass (*37*). It also allows to trace ^2^H into microbial neutral-lipid fatty acids (NLFAs), mainly consisting of triglycerides. NLFAs are considered as storage compounds found mainly in fungi and many prokaryotic species, they can account for a large C fraction of the total soil microbial biomass pool (*38*). While classically biomass growth is investigated by measuring cell replication, soil microbes can produce and accumulate large amount of storage compounds representing up to 46% of the total microbial biomass, especially in ‘stress’ situations (*29*). This trait can be particularly relevant in ecosystems experiencing large variations in C and nutrient supply (*39*) such as during drought conditions. Given the central roles of microbes in soil C cycling, methods that can quantify C invested in microbial growth (i.e., not only cell division but also storage compound synthesis) are strongly needed, particularly under current changing climate conditions.

Here, we investigated impacts of multiple climate change factors on the soil microbial community growth under field conditions. The overarching goal of this work was to quantify community level and group-specific growth rates of soil microbial communities under drought and potential interactions with simulated future climate conditions, i.e., the combined increase in temperature and atmospheric CO_2_ concentrations. We hypothesized that: (i) drought reduces microbial community growth rates, but the impacts of drought are reduced in a future climate, as both eCO_2_ and warming may stimulate microbial growth; (ii) fungal and bacterial growth display a different sensitivity to drought, with fungal growth being more resistant compared to more sensitive bacterial groups; and (iii) soil microbes increase the partitioning of C towards reserve compounds during drought. In a multifactorial climate change experiment, we manipulated atmospheric CO_2_ concentrations (+300ppm above ambient) and temperature (+3 °C above ambient) together (termed ‘future climate condition’ hereafter) and investigated how this condition altered the effects of drought compared to ambient controls (ambient, drought, future climate, future climate + drought). We applied ^2^H-PLFA-SIP in combination with a water vapor equilibration technique (*27*) to measure soil microbial growth rates under drought conditions and compared them to ^18^O-DNA-SIP obtained with the same technique (*40*). These methods are referred to as ^2^H-vapor-PLFA-SIP and ^18^O-vapor-DNA-SIP hereafter. This allowed us to link the coarse taxonomic resolution of microbial groups with a quantitative measure of growth rates and storage compound formation, and to compare PLFA-based to DNA-based growth rate estimates. We show that (i) soil microbial communities can maintain half of their growth rates under drought and quickly return to control levels after drought, indicating strong resilience; (ii) soil microbial CUE is insensitive to drought, but decreases under future climate conditions; (iii) fungi display remarkable resistance to drought, with no effect of drought on growth rates, and an increase in the investment into storage compounds, accounting for about three-fold the investment into growth. We further discuss the implications of microbial community responses to drought for soil biogeochemistry under future global change scenarios.

## Results

### Soil microbial communities maintain half of their growth rates during drought but recover quickly

We investigated community level physiological parameters (community level growth, respiration and CUE) with ^2^H-vapor-SIP (tracing ^2^H incorporation into fatty acids) and compared it to ^18^O-vapor-SIP (^18^O incorporation into DNA). Both methods estimate cell division rates and showed similar responses to the global change treatments, with drought significantly reducing mass specific growth rates by 48% (under both ambient and future climate; p=0.001, Table S1) when measuring ^2^H incorporation into PLFAs and by 57% to 63% (in ambient and future climate, respectively; p<0.001, Table S1) when tracking ^18^O into DNA (Fig. 1a and Fig. S1). Respiration rates responded similarly with a significant effect of drought (Fig. 1c and Fig. S1). Drought did not significantly change microbial community level CUE, but the future climate treatment caused a significant reduction in CUE (p=0.001; Table S1) during the recovery period only (Fig. 1b and Fig. S1). Mass specific growth rates and CUE calculated via ^2^H incorporation into PLFAs were consistently lower compared to ^18^O incorporation into DNA (Fig. 1), but the relative differences in response to treatments remained constant (supplementary Table S1). Mass specific growth rates and CUE significantly correlated between the two methods (Fig. S1), although ^2^H-vapor SIP based growth rates values were significantly (∼50%) lower.

**Figure 1.**
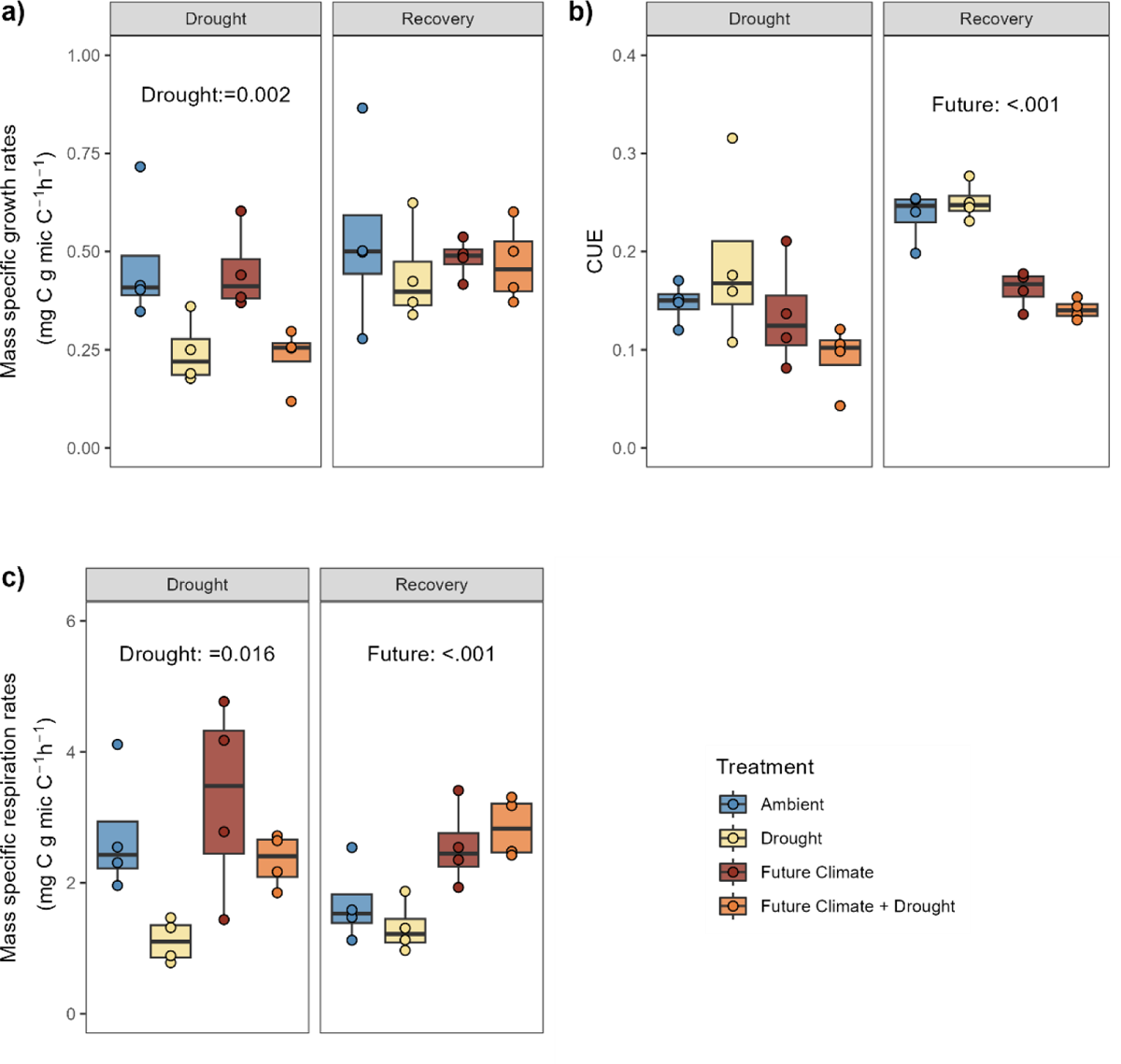
Soil microbial community-level mass specific growth rates, respiration and CUE during peak drought and recovery. Mass specific growth rates obtained via **(a)** ^2^H incorporation into PLFA **(b)** as well as CUE and **(c)** mass specific respiration rates, measured at two time points (‘Drought’ and ‘Recovery’). Significant differences (p<0.05) between treatments derived from linear mixed models are reported in the figure (the full report is provided in Table S1). Box centre line represents median, box indicates the upper and lower quartiles, whiskers the 1.5x interquartile range, and separated points represent potential outliers (n=4). Colour indicates treatment.

### The active microbial community changes in response to drought and future climate

Our ^2^H-vapor-PLFA-SIP approach to trace ^2^H into different PLFA biomarkers allowed not only tracking microbial activity (i.e., growth and CUE) at the community level, but also distinguishing the production of different fatty acids, indicative of the activity of different microbial groups, during drought. Drought, future climate and their interaction changed the active microbial community (displayed as principal component analysis, Fig. 2, Permanova results in Table S2) during drought. Drought separated the active community on PC1 (43%), dominated by a high incorporation of ^2^H into fungal markers during drought, relative to a low incorporation into gram-positive markers (Fig. 2). Future climate conditions favored the incorporation of ^2^H into gram negative and actinobacterial makers on PC2 (30.3%). Total PLFA abundance (not ^2^H incorporation) showed similar, but weaker patterns (Table S2 and Fig. S2). The impact of the preceding drought was maintained in the recovery period and significantly changed the relative ^2^H incorporation into PLFA biomarkers (Fig. 2 and Table S2) and relative PLFA abundances (Fig. S2 and Table S2). In the recovery period, future climate only significantly affected the relative abundance, but not the ^2^H incorporation into PLFAs (Fig. 2, Fig. S2 and Table S2).

**Figure 2.**
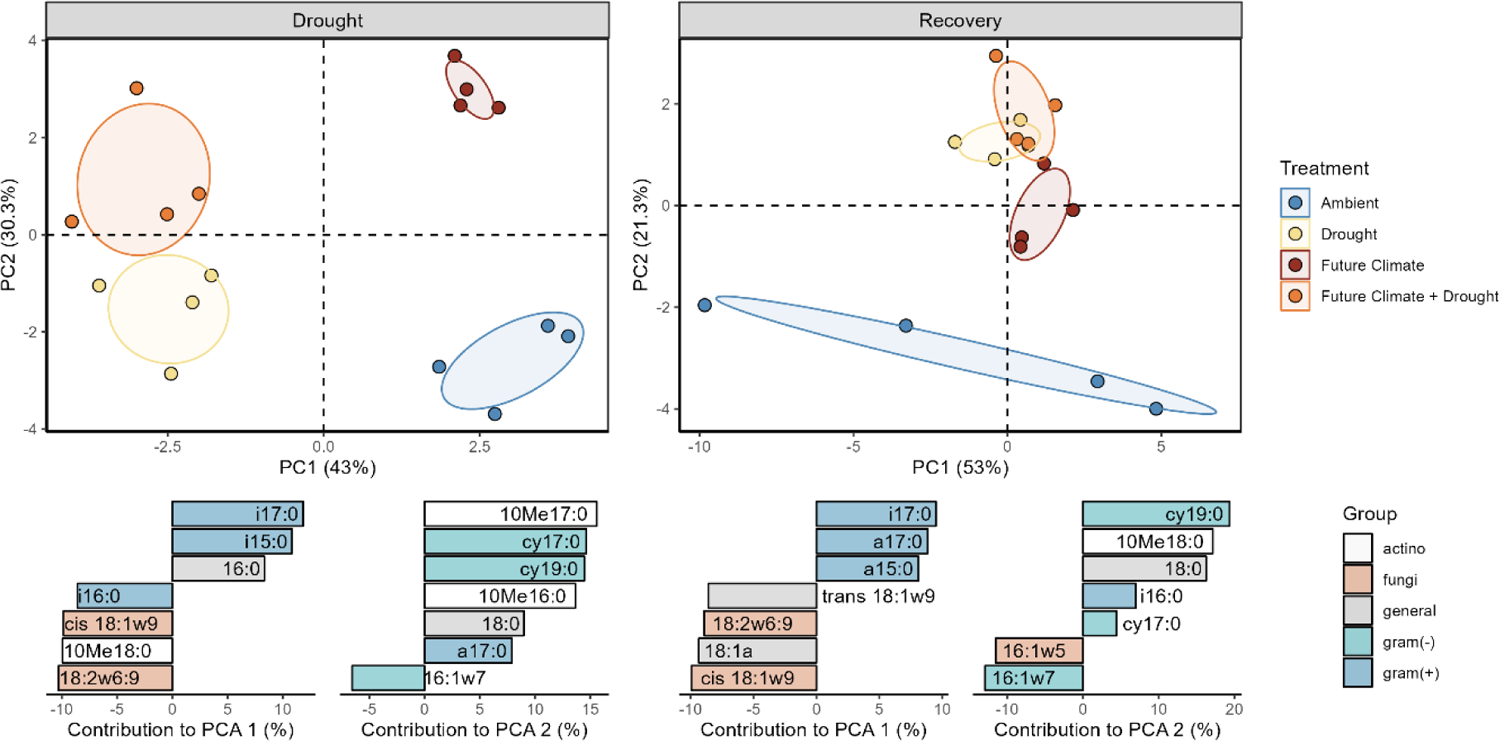
Multivariate analysis of ^2^H incorporation into individual PLFA biomarkers. PCA analysis of ^2^H incorporation into individual PLFA biomarkers during ‘Drought’ and ‘Recovery’ period. Bottom graphs represent the relative contribution (in %) of the top seven variables to the principal components (the positive or negative sign indicates the direction along the respective PCA axes) assigned to microbial groups (gram positive: blue; gram negative: light blue; fungi: orange; actinobacteria: white; general markers: grey; the arbuscular mycorrhiza fungal biomarker 16:1w5 is included in the fungi group in the multivariate analysis but not in other graphs displaying fungi). Permanova results are reported in Table S2. The sample size represents biologically independent samples (n=4). Ellipses represent the 95% confidence intervals. Colour indicates treatment.

### Fungal mass specific growth rates display remarkable drought resistance

Mass specific growth rates of gram positive, gram negative and actinobacteria markers (as well as general microbial markers) significantly decreased with drought, while fungal rates were not affected (Fig. 3, Fig. S3 and Table S3). At peak drought, gram-positive markers decreased on average by 49% in soils from ambient and 53% in future climate treatments, while gram negative markers decreased by 54% and 49%, respectively. The mass specific growth rate of fungi relative to bacteria was significantly increased under drought conditions by 84% and 173% in ambient and future climate, respectively (Fig. 3 and Table S3). The fungi to bacteria mass-specific growth returned to control values after rewetting (Fig. 3d). During the recovery period mass specific growth rates of all microbial groups became similar to levels measured in ambient soils (Fig. 3), with no significant effect of treatments.

**Figure 3.**
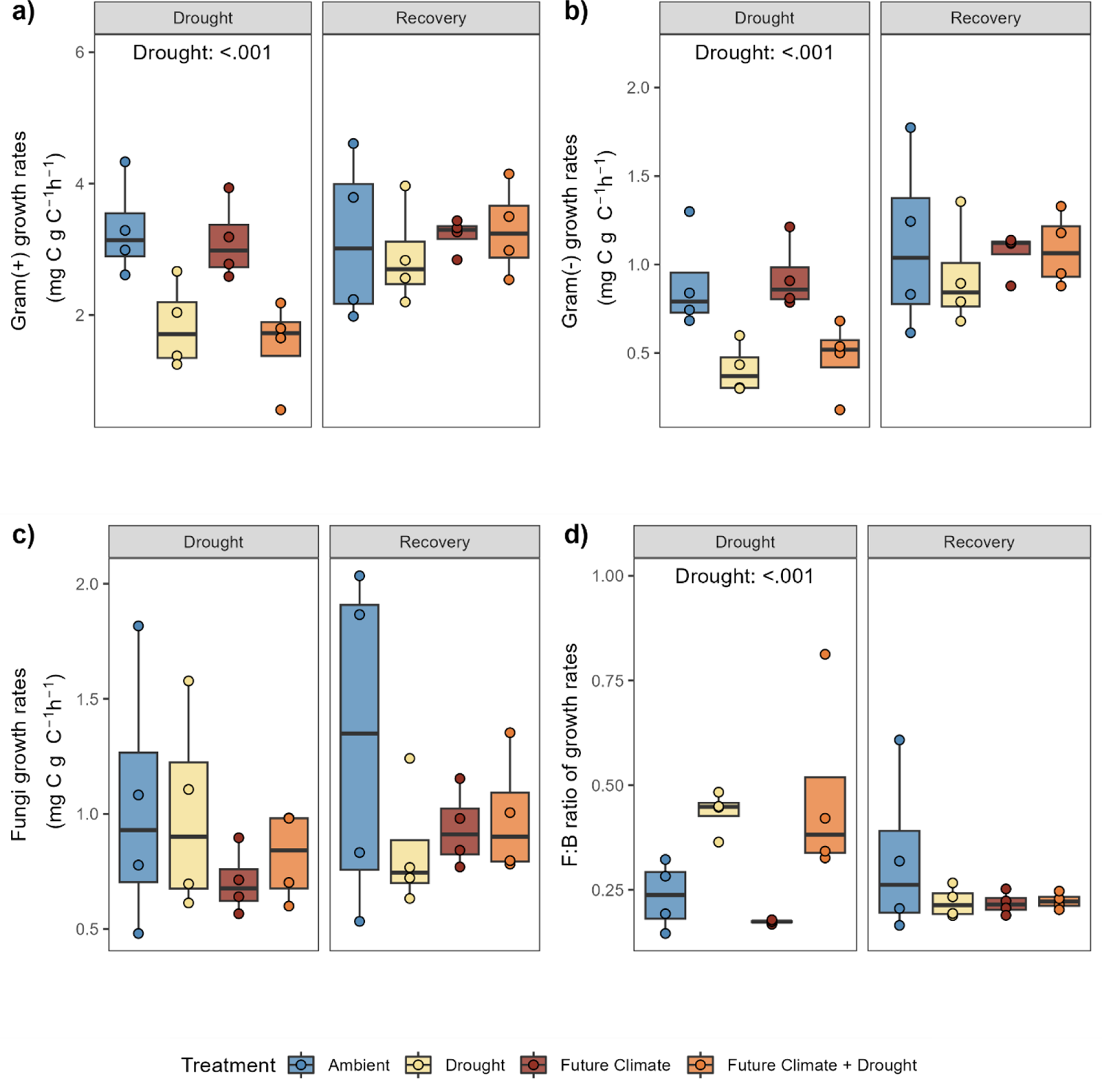
Mass-specific growth rates of different microbial groups. **a)** Gram-positive and **b)** Gram-negative bacterial markers and **c)** fungal markers, as well as **d)** the ratio of fungal to bacterial growth rates at peak drought and recovery (‘Drought’ and ‘Recovery’). Statistical results are reported for significant p-values (for a full report see Table S3). Box centre line represents median, box limits the upper and lower quartiles, whiskers the 1.5x interquartile range, while separated points represent potential outliers (n=4). Colour indicates treatment.

Mass specific growth rates correlated positively with respiration rates (Fig. S3). Actinobacteria and gram-negative bacteria displayed the strongest degree of correlation (r=0.87 and r=0.85, respectively), while fungi had the lowest degree of correlation (r=0.43) and the fungi to bacteria ratio correlated negatively with respiration rates (r=-0.38). As PLFAs are assumed to be a good indicator of viable biomass, we also compared results obtained with our SIP approach to results obtained by using absolute abundance of PLFA values. We found different results, with no overall significant effects except for future climate in gram positive and a significant effect of drought and rewetting on the fungi to bacteria ratio (supplementary Fig. S5). Correlation of abundance values with respiration rates were all positive, but displayed lower correlation coefficients, compared to correlations between mass specific growth rates and respiration rates (supplementary Fig. S6), and fungal abundance displayed the highest correlation coefficient with respiration rates.

### Fungi increase the synthesis of storage compounds during drought

NLFAs represent reserve compounds, which were detectable in fungi-associated fatty acid markers (18:1ω9cis and 18:2ω6,9), the marker 16:1ω7 (a biomarker for gram-negative bacteria), 16:1ω5 (arbuscular mycorrhizal fungi) and the general markers 16:0, 18:0 and 18:1ω9trans. We found that during drought the new production of fungal NLFAs was significantly higher compared to non-drought treated soils (Fig. 4a). The ratio of fungal NLFA to PLFA biomarkers (expressed as percentage) was on average 228 and 305% during drought, but only 22% under ambient and non-drought treated future climate (Fig. 4b). NLFA production rates decreased to ambient levels in the ‘Recovery’ period. Similar patterns were found for gram-negative bacterial markers, but with a smaller increase of the respective NLFA relative to PLFA marker production during drought (up to 6.3%; Fig. S6) and a faster decrease of NLFA production rates already after laboratory rewetting (Fig. S7 and Table S4).

**Figure 4.**
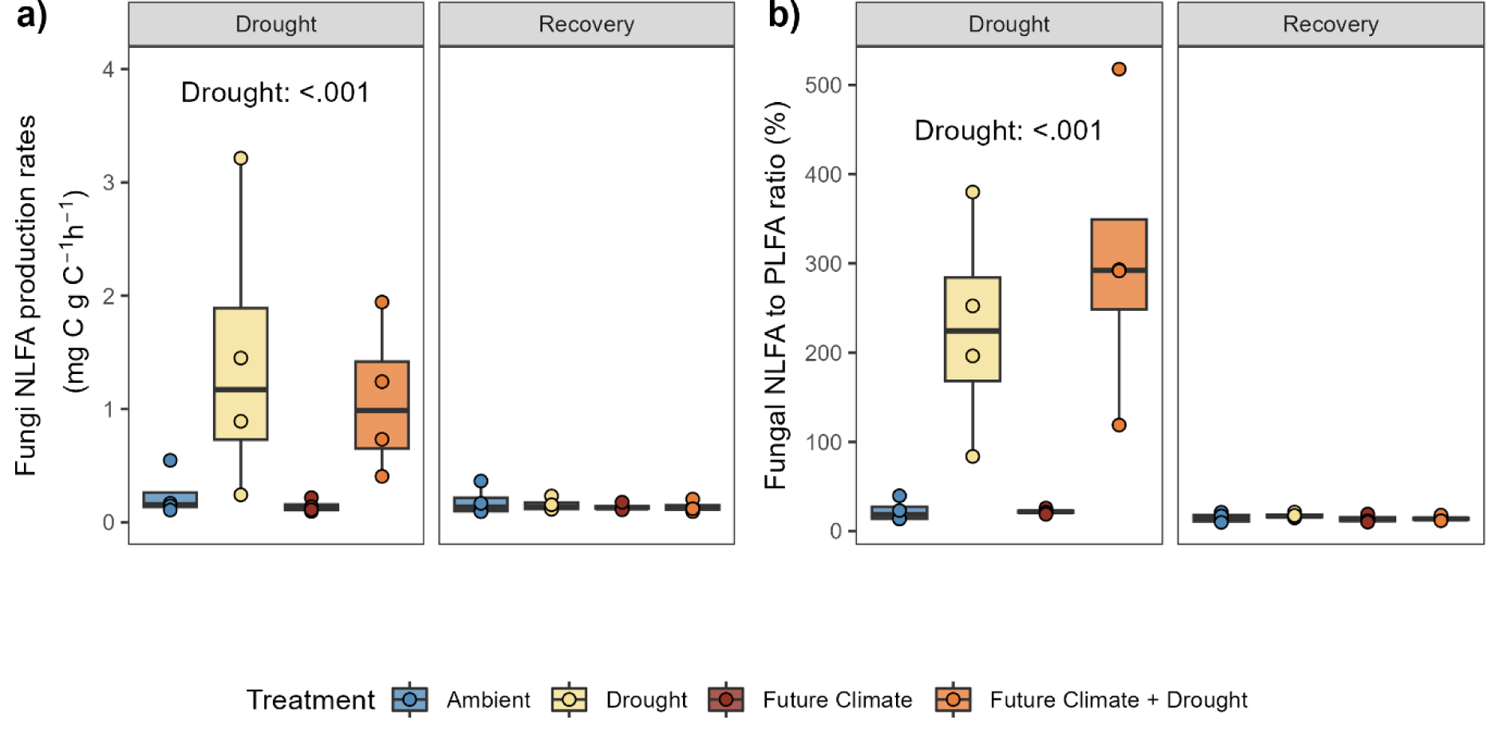
Fungal investment into storage compounds. **a)** Production of fungal specific NLFA during ‘Drought’ and ‘Recovery’ periods. **b)** NLFA to PLFA ratio (expressed as percentage) of fungal specific biomarkers production rates. Statistical results are reported for p-values <0.05 (for a full report see Table S4). Box centre line represents median, box limits the upper and lower quartiles, whiskers the 1.5x interquartile range, separated points indicate potential outliers. The sample size represents biologically independent samples (n=4). Colour indicates treatment.

## Discussion

Soil microbial biomass growth ultimately determines soil C and nutrient cycling and is thus a key variable in microbial ecology. Despite the significance of microbial growth dynamics for predicting the impacts of global environmental changes on soil biogeochemistry (*1*, *6–8*), there is still little direct evidence of responses of microbial growth rates under field conditions during drought and predicted future climate scenarios. Here we tackled the challenge by using a novel approach (^2^H-vapor-PLFA-SIP) that allows the concomitant estimation of community-level and group-specific C allocation to biomass growth and to storage product synthesis, relative to C mineralized to estimate community-level carbon use efficiency. We applied ^2^H-vapor-PLFA-SIP to responses of soil microbial communities to simulated drought and future climate conditions (eCO_2_ and warming) within a field experiment in a permanent grassland. We show that drought strongly reduces the overall growth of the soil microbial community, but at the same time, fungi display a remarkable resistance to drought independently of a future climate scenario.

Drought decreases the water filled pore space of soil, which constrains connectivity and carbon and nutrient resource diffusion, whereas in the remaining water filled pore space osmotic potential strongly decreases (*41*). In our experiment the simulated drought significantly reduced soil water content (by 73% on average) and in accordance with our hypothesis, community-level mass-specific growth rates decreased by half. This is in line with reports from previous lab incubations observing reduced microbial growth with lowering soil moisture (*42*, *43*). As in many previous studies substrates were applied dissolved in water, they could have triggered short-term rewetting responses and may have over-estimated microbial growth activity under very dry conditions (*42–48*). On the other hand, our results show that still 50% of the microbial community was remarkably resistant and remained active and producing PLFAs, even after two months of intense drought. Moreover, the microbial community showed a fast recovery after drought ended. These empirical results are extremely important, as data from drought field studies is largely unavailable and sustained microbial activity during drought and recovery conditions could have large effects on soil carbon storage (*18*, *49*). Finally, in contrast to our hypothesis future climate conditions neither buffered nor enforced drought and recovery effects on the microbial community level growth. While elevated CO2 and temperature can stimulate microbial growth (*47*), our results demonstrate the overriding effect of drought over other climate change factors.

Fostering a mechanistic understanding of soil C decomposition rates and soil functioning, requires innovative approaches to quantitatively link microbial community composition to soil processes (*50*– *52*). The use ^2^H-vapor-PLFA-SIP to estimate phospholipid fatty acids production rates allows to separate broad taxonomic groups across microbial kingdoms. Our results showed that the active community significantly differed in their composition between ambient conditions and future climate, as well as drought and rewetting (Fig. 2; Table S2). Our approach also demonstrates that the PLFA composition alone (only based on PLFA abundance) does not reliably describe the active fraction of the microbial community, especially during drought. While PLFAs can have a fast turnover time in soil (*53*) our results indicate that this might not hold true during drought conditions. This discrepancy between growth- and biomass-based assessment, highlights the importance of capturing growth rates with SIP approaches to understand microbial responses to environmental perturbations (*54*, *55*). The growth of different soil microbial groups (fungi, gram negative and gram positive) strongly differed in their sensitivity to drought, but was not affected by future climate conditions. Interestingly, both gram-positive and gram-negative bacteria similarly reduced growth during drought (Fig. 3), although gram positive being suggested to be drought resistant because of their cell wall structure (*56*, *57*). Fungal growth instead, showed an astonishing resistance to drought and exhibited unchanged growth rates after a two-month summer drought in the field. While fungi have often found to be more resistant in their community composition (*58*), based on biomass and DNA measurements (*15*, *58–62*) and accessing labile plant C in the rhizosphere during drought (*56*), to our knowledge this is the first study directly demonstrating that soil fungal growth in soil display complete resistant to field drought. Fungi can resist drought via their extensive filamentous-like body structure (*i.e.,* hyphae) that allows to reach water and nutrients even when drought reduces diffusivity (*15*, *63*, *64*). However, the lack of experimental approaches to eliminate potential biases caused by substrate and/or water addition has so far hindered the precise assessment of growth dynamics during drought conditions. Finally, our study also show bacterial growth was resilient and all groups showed a fast recovery after drought.

Microbial biomass growth is classically supposed to be directly represented by cell replication rates. However, microorganisms can allocate large amount of C to other processes, such as storage compounds, which can represent a large fraction of the total biomass (*29*). The ^2^H-vapor-PLFA-SIP method enabled us to additionally characterize the production rates of NLFAs (triglycerides), which represents major storage compounds in eukaryotic cells (*65*) and also in many bacterial species (*38*, *66*). In fungal cells triglycerides can account for up to 80% of the cell dry weight, in extreme situations (*67*), but their ecological role has not received much attention (*38*). Our results showed a large increase in storage compound synthesis during drought conditions, representing on average an increase of 5.8 (drought) and 4.8 times (future climate + drought) compared to non-drought conditions. NLFAs associated to fungi displayed the highest increase relative to the respective ambient conditions. Under non-drought conditions fungal storage compounds represented only around 22% of the investment in biomass (estimated by using the same PLFA biomarkers), which increased on average to 228% and 305% in drought and future climate + drought conditions, respectively, but production rates decreased fast during recovery and were indistinguishable from ambient controls. As it was shown that for arbuscular mycorrhizae, spores contain much more NLFAs that PLFAs, and NLFAs make about 20% of the total spore biomass (*68*). Hence, the fungal NLFA production observed in our study could also be an indication of spore formation and represent an important strategy for survival of extended periods of drought. Fungal investment into storage could represent an important strategy in relation to drought but it would have not been captured by using DNA-based methods. Moreover, ^2^H-vapor-PLFA-SIP could also be used to measure the production rate of other storage compounds during drought, such as polyhydroxybutyrate (PHB), which represent an important storage compound of bacteria (*69*).

Drought can have negative impacts on the soil carbon balance (*16*), especially if soil microorganisms can maintain higher respiration activity compared to primary productivity (*18*, *49*). Moreover, the increase in respiration and growth following rewetting of dry soils (‘Birch effect’; (*70*)) can lead to a decoupling of microbial anabolic and catabolic processes, and more C being used for respiration (*43*). Interestingly, in our study, microbial community level CUE, describing the partitioning of C directed to growth versus respiration (*63*, *71*), remained unaffected by drought and in the recovery period. This is in line with a previous study by Herron et al. (2009), where microbial CUE (determined by ^13^C-acetic acid vapor addition in a laboratory incubation) decreased only at very low moisture levels below 0.05 g H_2_O g^−1^ soil or −6.0 MPa (comparatively, drought treatments in our experiment reached on average 0.07 g H_2_O g^−1^ soil). In contrast to drought, we found a significant decrease in CUE by 43% and 38% (using ^2^H and ^18^O,respectively) caused by future climate scenario (the combination of eCO_2_ and warming), only during the recovery period, similarly to a previous study on the same site (*46*). Other studies also showed that CUE can decrease with warming (*72*). However, we previously showed that response to eCO_2_ and warming displayed nonlinear behavior and were strongly affected by seasonality (*46*), therefore these results should be interpreted with caution.

We targeted two processes linked to cell replication: fatty acid and DNA production. Community-level rates of ^2^H incorporation into phospholipid fatty acids correlated with ^18^O incorporation into DNA, and showed the same responses to drought and future climate conditions (Fig. S1). However, by using ^2^H incorporation into fatty acids, the calculated community level growth rate and CUE were lower compared to DNA based estimates. Specifically, per unit of PLFA fewer new PLFA was produced compared to the relative amount of newly formed DNA. Fatty acid ^2^H values reflect the isotopic composition of the source water (*73*). The amount of ^2^H derived from added labelled water can be considered as a combination of the mole fraction of water derived H and associated net ^2^H isotope fraction (*74*). Therefore, to accurately estimate microbial fatty acid production the physiological parameter ‘water hydrogen assimilation constant (a_w_)’ should be applied (*74*, *75*), obtained from the regression of the ^2^H isotopic composition of fatty acids and water source. In a recent survey it was demonstrated that a_w_ can range from values close to 0.1 up to values close to 1 (*76*). In complex communities, such as soil, the assimilation constant is difficult to estimate, but assuming all microbes are heterotrophs, this value is on average 0.71 (*35*), which we also applied in this study. However, it is important to note that using a different aw factor could lead to strong changes in the final quantification. For example, if we had used a value of 0.3, we would have generated almost identical PLFA and DNA-based growth rate estimates. Nevertheless, relative differences between treatments in our experiment are independent of the used value.

Understanding terrestrial ecosystem response to climate change events, such as drought, requires innovative approaches to quantify microbial growth and soil carbon cycling. Using ^2^H-vapor-PLFA-SIP allows the concomitant measurement of community level physiology and the direct quantification of growth and storage product synthesis of different microbial groups. Our results demonstrate that drought exerts a major control on microbial catabolic and anabolic processes. Despite the strong effects on microbial growth and respiration, soil microbial communities are able to maintain the balance between these processes, with stable microbial CUE values across drought and rewetting. Most notably, we could unequivocally demonstrate that soil fungi remain active during a two-months field drought event. During drought fungi invest large amounts of carbon into intracellular storage compounds, a strategy that could be connected to fungal resistance and resilience to drought. Accounting for different physiological strategies of growing soil microbes during global change, i.e., cell division versus storage compound synthesis, can foster our understanding of soil microbial contribution to the C cycle and ecosystem feedback to climate change.

## Methods

### Site description, experimental layout and sample collection

Samples were collected from a managed montane grassland in the Austrian Alps, Styria, Austria (47°29’38’’N, 14°06’03’’E) as part of a multifactorial climate change experiment (‘ClimGrass’) located at the Agricultural Research and Education Center (AREC) in Raumberg-Gumpenstein. The site is characterized by a mean annual temperature of 8.5 ⁰C and a mean annual precipitation of 1077 mm. According to the WRB-system (*77*) the soil is classified as Dystric Cambisol (arenic, humic) with a loamy sand texture and a pH-value of ∼5.5. Before establishment of the ‘ClimGrass’ experiment, a typical grassland mixture was sown in an area of homogeneous soils in 2007 (species list is descried in the supplementary text). The ClimGrass project entails 54 plots with a combined warming and Free-Air-Carbon dioxide-Enrichment (T-FACE) setup, put into full operation in 2014 to manipulate temperature and CO_2_ at three levels each (*46*, *78*, *79*). Fully automated rainout shelters were installed above half of the ambient and above half of the combined +3.0 °C and +300 ppm CO_2_ (i.e. ‘Future Climate’) plots. All plots are harvested (plant biomass) three times a year (spring, summer and autumn) and receive identical rates of mineral fertilizer, applied in three batches giving a total load of 90 kg N, 65 kg P, 170 kg K; per hectare and year.

For this experiment, we chose 16 plots representing four different treatments in a full factorial design (n=4 per treatment, respectively): ambient (‘ambient’), drought (‘drought’), eCO_2_ and elevated temperature combined (+300ppm +3°C; ‘future climate’), and future climate with drought (‘future climate + drought’). The drought period was simulated in the field between June 17^th^ 2020 until August 3^rd^ 2020 by excluding all naturally occurring precipitation. The drought plots then received a scheduled rewetting with 40 mm of previously collected rainwater on August 3^rd^ 2020, after which the automatic rain-out shelters were switched off and the plots were used to investigate the effects of drought recovery.

We collected soil samples towards the end of the drought period (29^th^ of July) and two days after the rewetting event (5^th^ of August). Hereafter, we refer with *‘Drought’* to the samples that were collected at the end of the severe drought period, and with *‘Recovery’* to the soil samples that were collected after rewetting. From each experimental plot, three soil samples were collected using a soil corer with a diameter of 2 cm and a length of 10 cm. Per plot, samples were pooled, fresh masses weighed, sieved to 2 mm and fine roots were removed. Aliquots of fresh sieved soil were weighed and dried (105 °C, 48 h) to calculate soil water content. Further fresh soil aliquots were used for determining microbial biomass based on the chloroform fumigation extraction method described by (*80*), as well as setting up the laboratory incubations (see below).

### Laboratory incubation set-up: soil microbial activity using ^2^H-vapor-SIP compared to ^18^O-vapor-SIP

We tested the applicability of ^2^H incorporation into phospho- and neutral lipid fatty acids (PLFA and NLFA) to determine soil microbial group specific growth and buildup of storage compounds under drought and future climate change conditions. For applying labelling soil water with ^2^H we used the water vapor equilibration method as described by Canarini et al. (2020) for ^18^O incorporation into DNA. For clarity, we will refer to the ^2^H-tracing approach as ^2^H-vapor-SIP. In addition, we compared the PLFA-based (^2^H-vapor-SIP) to DNA-based (^18^O-vapor-SIP) estimates for microbial growth and carbon use efficiency (CUE_PLFA_ vs. CUE_DNA_). The short-term laboratory incubations were started within 24 h after sample collection.

We used a similar experimental set up as previously described in details (*27*), but adjusted soil sample size based on the amount of soil needed for extracting sufficient amounts of fatty acids or DNA, respectively: we incubated around 800 mg (in duplicates) to trace ^2^H-labelled water into PLFAs and NLFAs and around 400 mg of fresh soil to trace ^18^O labelled water into DNA. Soil samples were weighed into 1.2 ml plastic vials and inserted in 27 ml glass headspace vial, which were closed air-tight with rubber septa. For both assays (^2^H-vapor-SIP and ^18^O-vapor-SIP) the labelled water was applied at the bottom of the glass headspace vial with no direct contact to the soil. The amount of water added was calculated based on the amount of water that would increase the soil water to 60% of their respective water holding capacity to aim for an approximate enrichment of the soil water of around 20 at% ^2^H and 20 at% ^18^O, by the end of the incubation.

Samples were incubated at their respective field temperatures at the time of sample collection (Ambient, Drought: 20°C; Future Climate, Future Climate + Drought: 23°C) for 48 h. For both assays CO_2_ concentrations in the glass headspace vials were determined at the beginning and end (after 48 h) of the incubation to calculate soil respiration rates (see details below). At the end of the incubation the soil samples were removed from the headspace vials, closed, and shock frozen in liquid N_2_, then kept at −20 °C until further analyses.

### Temporal dynamics of ^2^H and ^18^O equilibration with soil water

At the ‘*Drought*’ collection we randomly selected several samples (two for each treatment) for both ^2^H and ^18^O incubations. From this we collected aliquots of the remaining isotopically labelled water (^2^H or ^18^O) from the bottom of the vial after 3, 6 and 16 h of incubation. In addition, we collected the water at the end of the 48 h incubation, to calculate the incorporation of ^2^H and ^18^O into soil water as described in Canarini et al. (2020) (for details see Supplementary text and Fig. S8 and S9).

The collected ^2^H-labelled water was analyzed using platinum catalyzed equilibration of ^2^H in H _2_O with H_2_ gas by a Gasbench II headspace sampler connected to a Delta V Advantage isotope ratio mass spectrometer (Thermo Fisher, Bremen, Germany). The sampled ^18^O-labelled water was analyzed through equilibration of ^18^O in H _2_O with CO_2_ by a Gasbench II headspace sampler connected to a Delta V Advantage isotope ratio mass spectrometer (Thermo Fisher, Bremen, Germany). Both ^2^H and ^18^O values of the collected water samples were calibrated against isotope calibration curves using water with known isotopic values. We calculated the ^18^O and ^2^H soil water enrichment via measuring loss of enriched isotope in the added source water (Canarini et al. 2020). The ^2^H incorporation showed slower equilibration curves but reached comparable isotopic enrichment to the ^18^O method after 48 h (Fig. S8 and S9). Additionally, we verified that ^2^H does not inhibit microbial activity. We measured respiration in both labelled and natural abundance samples and tested with a pair t-test if respiration rates were altered by the introduced ^2^H label. Results show no significant difference (t=1.992; p=0.0534; Fig. S10a) and that data aligns around the 1:1 line of respiration rates of labelled and natural abundance samples (Fig. S10b).

### Microbial growth, respiration and CUE determined via ^2^H-vapor-SIP

Soil microbial growth, microbial respiration and microbial carbon use efficiency (CUE_PFLA_) were determined based on the incorporation of ^2^H from soil water into PLFAs. Additionally, we also calculated investment into lipid storage compounds via the incorporation of ^2^H from soil water into NLFAs (extracted as described below). In fatty acids the hydrocarbon skeleton consists of non-exchangeable C-H bonds which allows to use ^2^H-incorporation as an indicator for biological activity (*81*). Fatty acid biosynthesis and subsequently also ^2^H incorporation combine fatty acid production related to membrane growth, but also membrane repair; therefore, specific growth rates need to be considered accordingly. Microbial respiration was determined by measuring the CO_2_ concentration in the headspace vial right after the application of ^2^H enriched water and 48 h after the incubation using an infrared gas analyzer (EGM4, PP systems).

PLFAs and NLFAs were extracted from freeze-dried soil samples with a modified high throughput method (*66*). Total lipids were extracted from around 1 g of soil using a chloroform:methanol:citric acid buffer mixture and fractionated by solid-phase extraction on silica columns. The neutral lipid fatty acid (NLFA) fraction was collected by eluting samples with chloroform (containing 2% ethanol as recommended in (*82*), subsequently, the PLFA fraction was collected by eluting columns with a 5:5:1 chloroform:methanol:water mixture. After an internal standard (fatty acid methyl ester: 19:0) was added, NLFAs and PLFAs were converted to fatty acid methyl esters by transesterification. Samples were analyzed for H quantity and ^2^H incorporation using a Trace GC Ultra connected by a GC-IsoLink to a Delta V Advantage Mass Spectrometer (all Thermo Fisher Scientific). Samples were injected in splitless mode (injector temperature 300 °C) and separated on a DB5 column (60 m × 0.25 mm × 0.25 μm; Agilent, Vienna, Austria) with 1.2 ml He min^−1^ as the carrier gas (GC program: 1 min at 80 °C, 30°C min^−1^ to 150 °C, 1 min at 150 °C, 4 °C min^−1^ to 230 °C, 30 min at 230°C, 30 °C min^−1^ to 280 °C and 19 min at 280 °C). The H3 factor (correction for abundance of the trihydrogen ion H_3_^+^) was defined based on multiple injections of reference hydrogen gas before each sequence, and was very stable over time (3.64 ± 0.05 ppm/nA). FAMEs were identified using mixtures of bacterial and fungal FAMEs (Bacterial Acid Methyl Ester CP Mixture (Matreya LLC, State College, PA, USA) and 37 Comp. FAME Mix (Supelco, Bellefonte, PA, USA)), and quantified against the internal standard (19:0). Sample δ^2^H values were normalized to the VSMOW scale using the slope and intercept of measured and known values of isotopic standards. These were individual and fatty acid mixes of USGS70 and USGS72 (Arndt Schimmelmann, Indiana University) measured before and after each measurement run, as described above, and ranging from δ^2^H-183‰ to 384‰. Offsets between measured and known values for these standards were used to correct for any drift over the course of the sequence or any isotope effects associated with peak area or retention time. The standard deviation for these standards averaged 4‰ and the average offset from their known values was 27‰. Memory effect corrections were not applied; the maximal potential offset ranges around 4% (*83*) and analysis of methane reference standards that follow analytical peaks suggested that memory effect should be at around 0.2% (*75*).

We used the markers 18:1ω9cis and 18:2ω6,9 to estimate fungal biomass (*84*). For gram-positive bacteria we used the sum of i15:0, a15:0, i16:0, i17:0 and a17:0 markers and of the actinobacteria markers (10Me16:0, 10Me17:0 and 10Me18:0), for gram-negative bacteria we used the markers 16:1ω7, cy17:0 and cy19:0 (*85*). Fungi to bacteria ratio was calculated as the ratio between the fungal biomass and the sum of gram positive and gram negative. We used the markers 16:1ω5 as indicative of arbuscular mycorrhizae fungi, but kept separated from the other fungal biomarkers (except for the multivariate analysis). The remaining markers including 18:1ω9trans, 16:0, 17:0, and 18:0 cannot be exclusively assigned to bacterial or fungi and markers with double bond position which could not be chromatographically resolved, were assigned to the general PLFA marker group (*85*).

We calculated the newly produced C in each fatty acid (FA C_produced_) and expressed as ng C g^-1^ soil dry weight, using the enrichment of ^2^H as:

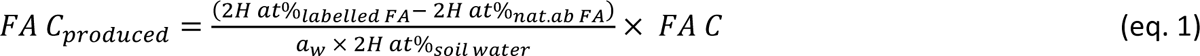

with ^2^H at%_labelled FA_ describing the atom% ^2^H of a specific fatty acid, ^2^H at%_nat.ab. FA_ representing the natural ^2^H abundance of the same fatty acid (sample taken from the same plot and incubated with natural abundance water), at%_soil water_ indicating the ^2^H atom% in the soil water estimated by the equilibration curves described in Supplementary methods. The factor a_w_ represents the assimilation efficiency of deuterium into fatty acids and it has been estimated at a value of 0.71 for heterotrophs (*35*, *74*). ‘FA C’ is the C concentration of the respective fatty acid (average of labelled and natural abundance sample) calculated relative to the known amount of C of the internal standard 19:0.

For each sample we calculated the total soil microbial community C production as the sum of FA C*_produced_* for PLFAs (PLFA C*_produced_*) or NLFAs (NLFA C*_produced_*), respectively. PLFA C_produced_ can be used to calculate microbial biomass growth rates (Growth_PLFA_ in ng C g^-1^ dw soil h^-1^), using soil microbial biomass C determined by chloroform fumigation extraction (*80*) as follows:

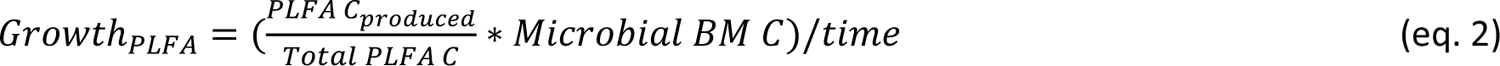

by multiplying the fraction of PLFA-C*_produced_* per total PLFA C (i.e. the sum of all FA C per sample, Total PLFA C) with the amount of microbial biomass C (Microbial BM C in µg g^-1^ dw soil) of each soil and divided by the incubation time. The amount of C taken up by the microbial community (C_Uptake PLFA_) was estimated as

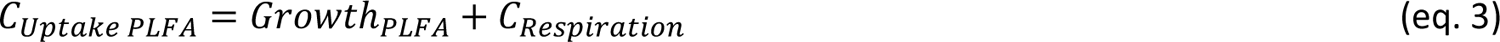

where Growth_PLFA_ is the C allocated to biomass production (ng C g^-1^ dw soil h^-1^), and C_Respiration_ is the C allocated to the production of CO_2_ (ng C g^-1^ dw soil h^-1^). In addition, we calculated mass-specific growth rates for the Growth_PLFA_ (PLFA-MS Growth rates) by dividing the values by Microbial BM C (final unit: mg C g^-1^ mic C h^-1^), and also for each microbial group separately, by summing up FA-C*_produced_* for each group using the assignment of FA described above (for both PLFA and NLFA separately), divided by the respective FA C and incubation time (mg C g^-1^ C h^-1^). For NLFA we also calculated the percentage of NLFA biomarker produced relative to the same PLFA biomarker and grouped by microbial group as per the assignment described above. Microbial CUE_PLFA_ was then calculated by the following equation (*63*, *71*):

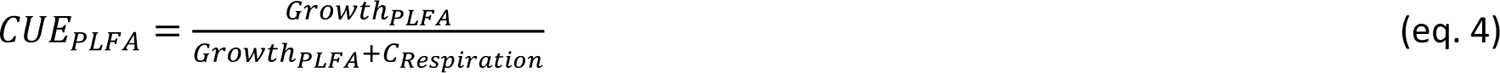

### Microbial growth respiration and CUE determined via ^18^O-vapor-SIP

As comparison to PLFA based microbial growth and CUE, we also determined DNA based growth (Growth_DNA_) and carbon use efficiency (CUE_DNA_) based on the incorporation of ^18^O from soil water into genomic DNA via ^18^O-vapor-SIP as described earlier (*27*). Microbial respiration was determined by measuring the CO_2_ concentration in the glass headspace vial as described above. DNA was extracted using a DNA extraction kit (FastDNA™ SPIN Kit for Soil, MP Biomedicals). DNA concentration of each extract was determined fluorimetrically following the Picogreen assay (Quant-iT™ PicoGreen® dsDNA Reagent, Life Technologies). Subsequently, the ^18^O enrichment and the total O content of the purified DNA fractions were measured using a Thermochemical elemental analyzer (TC/EA Thermo Fisher) coupled via a Conflo III open split system (Thermo Fisher) to an isotope ratio mass spectrometer (Delta V Advantage, Thermo Fisher). The amount of DNA produced over the 48 h of incubation can be calculated using the following formula as also described in Canarini et al (2020):

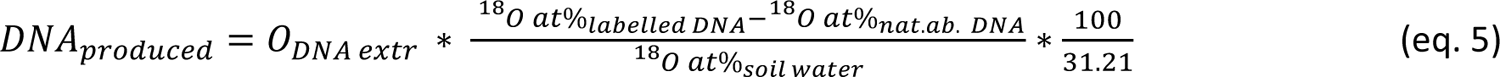

where *O_DNA extr_* is the total amount of oxygen in the DNA extract, *^18^O at%_DNA L_* and *^18^O at%_DNA n.a._* are the ^18^O enrichments of labeled and unlabeled DNA extracts, respectively, *^18^O at%_soil water_* is the ^18^O enrichment of the soil water (calculated as average ^18^O enrichment in soil water from the equilibration curves described in Supplementary methods), multiplied by the fraction of the average oxygen content in DNA (31.21%). We here used the amount of DNA_produced_ (µg g^-1^ soil dry weight) to calculate microbial growth in units of C by multiplying it with the ratio of microbial biomass C (determined by chloroform fumigation extraction (*80*) to DNA content of each soil sample:

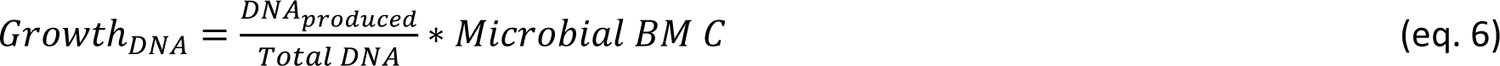

Similarly, the amount of C taken up by the microbial community based on ^18^O-vapor-SIP into DNA (C_Uptake_) is estimated as:

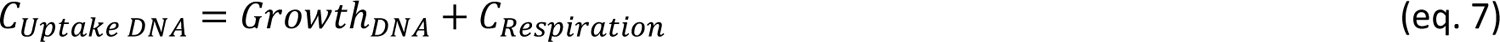

where Growth_DNA_ is the C allocated to biomass production, and C_Respiration_ is the C allocated to the production of CO_2_.). In addition, we calculated mass-specific growth rates for the Growth_DNA_ (DNA-MS Growth rates) by dividing the values by Microbial BM C Microbial CUE_DNA_ was then calculated as described above:

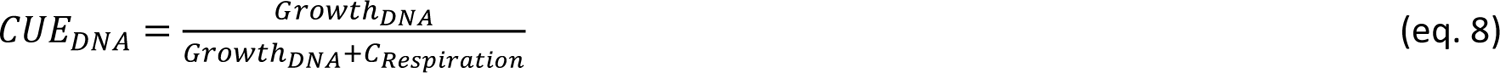

### Statistical analyses

Statistical differences in community level mass specific growth rates, microbial abundance, CUE and NLFA production rates were assessed by a linear mixed effect model separated for peak drought and rewetting sampling, using the package ‘nlme’ (*86*). For the harvest ‘Rewet’ we compared the rewetted samples to the non-drought controls of the harvest ‘Drought’. In all the models we tested for the effects of drought (or rewet), future climate and their interaction. Plot number was used as a random factor to account for plot variability. Differences in respiration between samples where ^2^H was added compared to natural abundance water were evaluated via two-tailed paired t-test, via the R function ‘t.test’. Correlations between CUE and mass specific growth for the method comparison were assessed via Pearson’s correlation via the ‘stat_cor’ function. Model assumptions were inspected visually, and values were log-transformed when assumptions were not met. Principal component analyses of relative PLFA abundances and growth rates were assessed via the function ‘PCA’ of the package ‘FactoMineR’ (*87*) and scores were normalized by setting scale = TRUE. Statistical significance of relative PLFA abundances and growth rates between treatments was assessed via PERMANOVA with the function ‘adonis’ of the ‘vegan’ package (*88*) using Euclidean distances. For both PCA and PERMANOVA data was split, with ‘Drought’ and ‘Rewet’ harvest analyzed together and ‘Recovery’ separately. Plots were generated via the package ‘ggplot2’ (*89*). Statistical analyses were performed in R 3.6.3 (*90*).

## Acknowledgments

We thank Ludwig Seidl for technical support with GC-IRMS measurements. We would like to thank the team of the Austrian Research and Education Centre Raumberg-Gumpenstein (HBLFA) for their support during the sampling campaign and for the provision of the experimental site, which was supported by the DaFNE project ClimGrassEco (101067) and funding of the Earth System Sciences research program of the Austrian Academy of Sciences (ÖAW project ClimGrassHydro) obtained by MB. LF acknowledges the European Union’s Horizon 2020 research and innovation program under the Marie Sklodovska-Curie grant agreement no. 847693 (REWIRE). DM was financially supported by FutureArctic, a European Union’s Horizon 2020 research and innovation program under the Marie Skłodowska-Curie Actions (grant no. 813114).

## Data availability

Data will be made available upon acceptance for publication on a public repository.

