## Supplementary file for "Soil fungi remain active and invest in storage compounds during drought independent of future climate conditions"

**This file includes:**

Supplementary Text
Figures S1 to S9
Tables S1 to S4

**Supplementary Text**

Site description

Before establishment of the global change experiment (ClimGrass), a typical grassland mixture was sown in an area of homogeneous soils in 2007, comprising the grass species *Arrhenatherum elatius* L., *Dactylis glomerata* L., *Poa pratensis* L., *Alopecurus pratensis* L., *Festuca rubra* L., *Trisetum flavescens* L., *Lolium perenne* L., *Phleum pratense* L. and *Festuca pratensis* L., and the legumes *Lotus corniculatus* L. and *Trifolium repens* L..

Equilibration curves of $^2H$ and $^{18}O$

As the $^2H$ and $^{18}O$ enrichment of the soil water is used to calculate PLFA and DNA production and ultimately microbial growth, the average $^2H$ and $^{18}O$ enrichment of soil water needs to be calculated across incubation time (48 hours) to account for the temporal dynamics of isotope equilibration of soil water. To do so, we used the same approach as described in Canarini *et al.*, (2020). This was achieved by collecting the water left at the bottom of the headspace vials and used to analyze its $^2H$ and $^{18}O$ enrichment in a subset of samples (first harvest 'Drought'). This was done at the end of the incubation period for all the samples (48 hours), and several times across the incubation period. We selected two random replicates per treatment, repeated the same set up as described in Materials and Methods section 2.2 and used these samples to collect the water left at the bottom of the headspace after 3, 6 and 16 hours after the start of the incubation. The curves obtained, representing the equilibration of the $^2H$ and $^{18}O$ labelled external water via the vapor phase with soil water, was fitted with a negative exponential function (eq. 1) as described in Ingraham and Criss (1993):

$$^{2}H\ at\%_{soil\ water} = \ ^{2}H\ at\%_{48} + \left(\ ^{2}H\ at\%_{in} - \ ^{2}H\ at\%_{48}\right) * \left(e^{-bt}\right) \qquad \text{(eq. 1)}$$

where $^{2}H\ at\%_{48}$ and $^{2}H\ at\%_{in}$ represent the $^2H$ atom % (or $^{18}O$) of the water after 48 hours incubation and at time point 0, while *b* represents a soil specific coefficient that was generated by fitting the *nls()* function in R. We then used the values obtained from this procedure for $^{2}H\ at\%_{48}$and *b*, to generate a prediction of the soil water isotopic enrichment as was described for $^{18}O$ demonstrated in Canarini et al., (2020). Finally we calculated the integral of this function by using the function *integrate()* of the R package "pracma" between time 0 and 48 hours. This integral was divided by 48 hours to generate an average isotopic enrichment (the term $at\%_{soil\ water}$ in equation 1 and the term $^{2}H\ at\%_{soil\ water}$ in equation 5 of the Materials and Methods section) of soil water for each treatment (non-droughted values were used for the treatments in the 'Rewet' and 'Recovery' period).

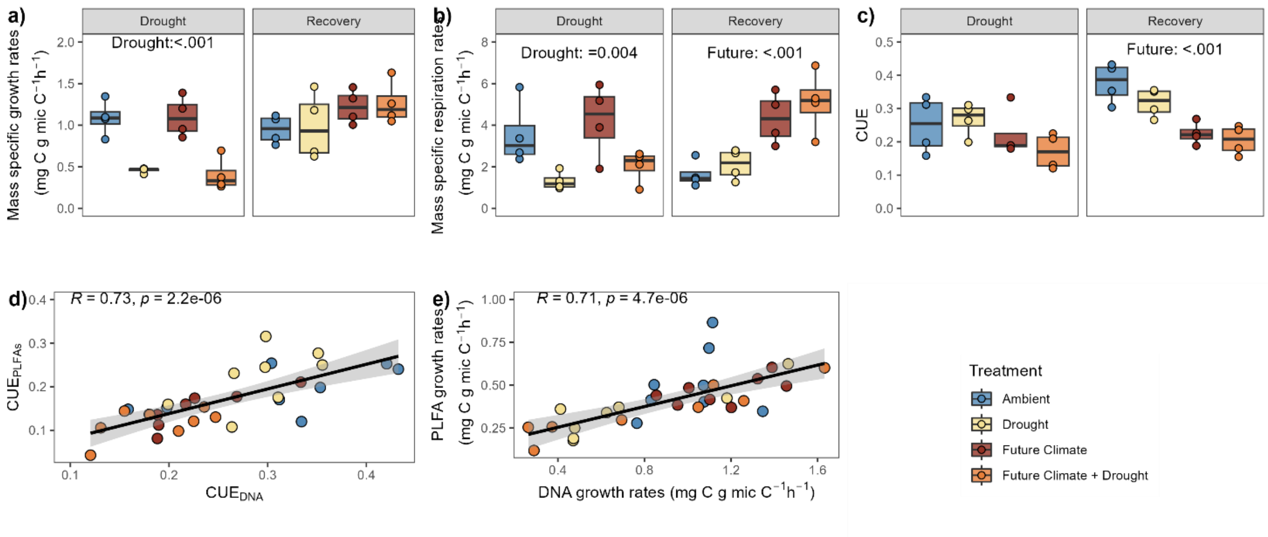


**Fig. S1.**

**Soil microbial community-level mass specific growth rates, respiration and CUE measured via[18]O incorporation into DNA and correlation with [2]H incorporation into PLFA**. Mass specific growth rates obtained via $^{18}O$ incorporation into DNA (a), as well as respiration rates (b) and CUE (c), measured at two time points (‘Drought’ and ‘Recovery’). Significant differences ($p<0.05$) between treatments derived from linear mixed models are reported in the figure (the full report is provided in Table S1). Box centre line represents median, box indicates the upper and lower quartiles, whiskers the 1.5x interquartile range, and separated points represent potential outliers (n=4). The bottom panels compare d) PLFA and DNA-based microbial community carbon use efficiency (CUE)  and f) mass-specific growth rates, obtained by the two methods. Pearson’s correlation coefficient R (and p-values) are reported on the graph. Colour indicates treatment.

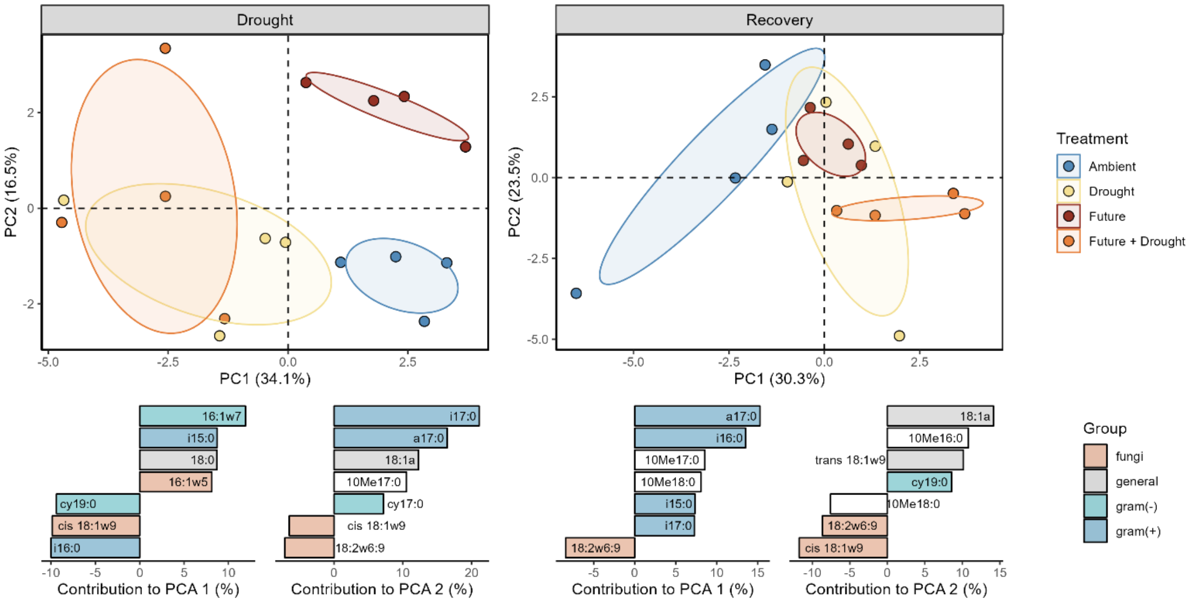


**Fig. S2.**

**Microbial community composition based on the relative abundance of individual PLFA biomarkers displayed as PCA** during 'Drought' and 'Recovery' period. Bottom graphs represent the relative contribution (in percentage) of the top 7 variables to the principal components (absolute values represent the relative contribution while the positive or negative sign represents the direction along PCA axes) divided by microbial group (Gram positive = blue; gram negative = light blue; general = grey; fungi = orange; actinobacteria = white; note that the arbuscular mycorrhizae fungal biomarker 16:1w5 is included into the fungal group in the multivariate analysis but not in other graphs displaying fungi). Permanova statistical results are reported in Table S2. The sample size 'n' represents biologically independent samples (n=4). Ellipses represent the 95% confidence intervals. Colour indicates treatment.

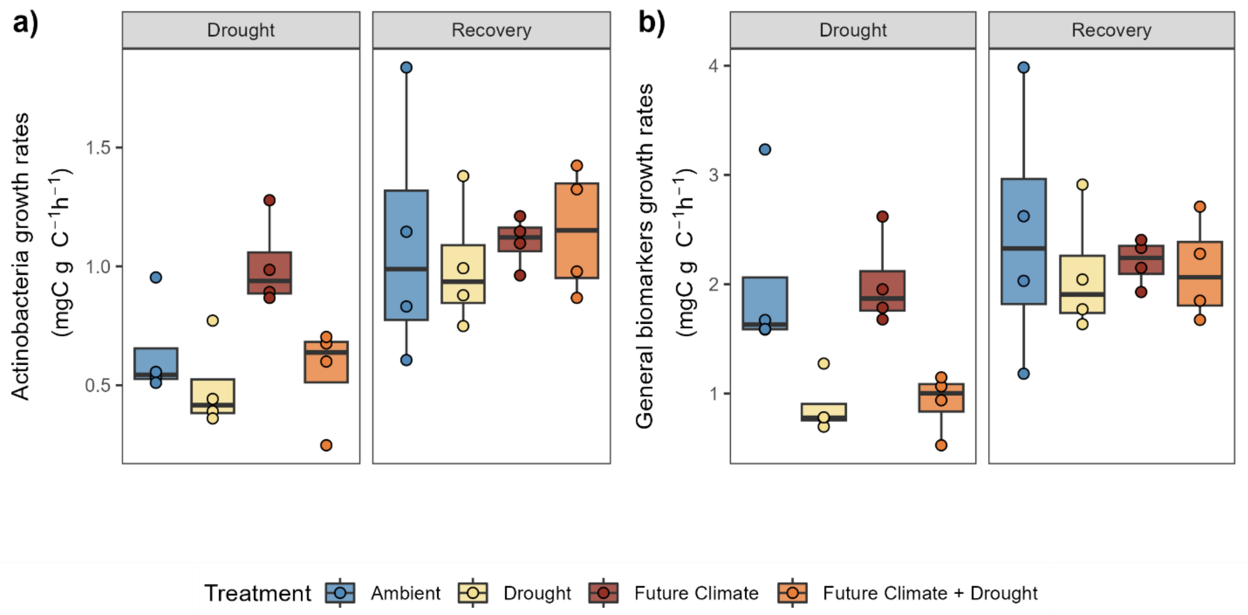


**Fig. S3.**

**Mass-specific growth rates** of a) actinobacteria and b) general biomarkers, divided by the two time points measured (Drought and Recovery). Statistical results are reported in Table S3. Box centre line represents median, box limits the upper and lower quartiles, whiskers the 1.5x interquartile range, while separated points represent potential outliers. The sample size 'n' represents biologically independent samples (n=4). For all the graphs colour indicates treatment (Ambient = blue, Drought = yellow, Future Climate = red, Future Climate + Drought = orange).

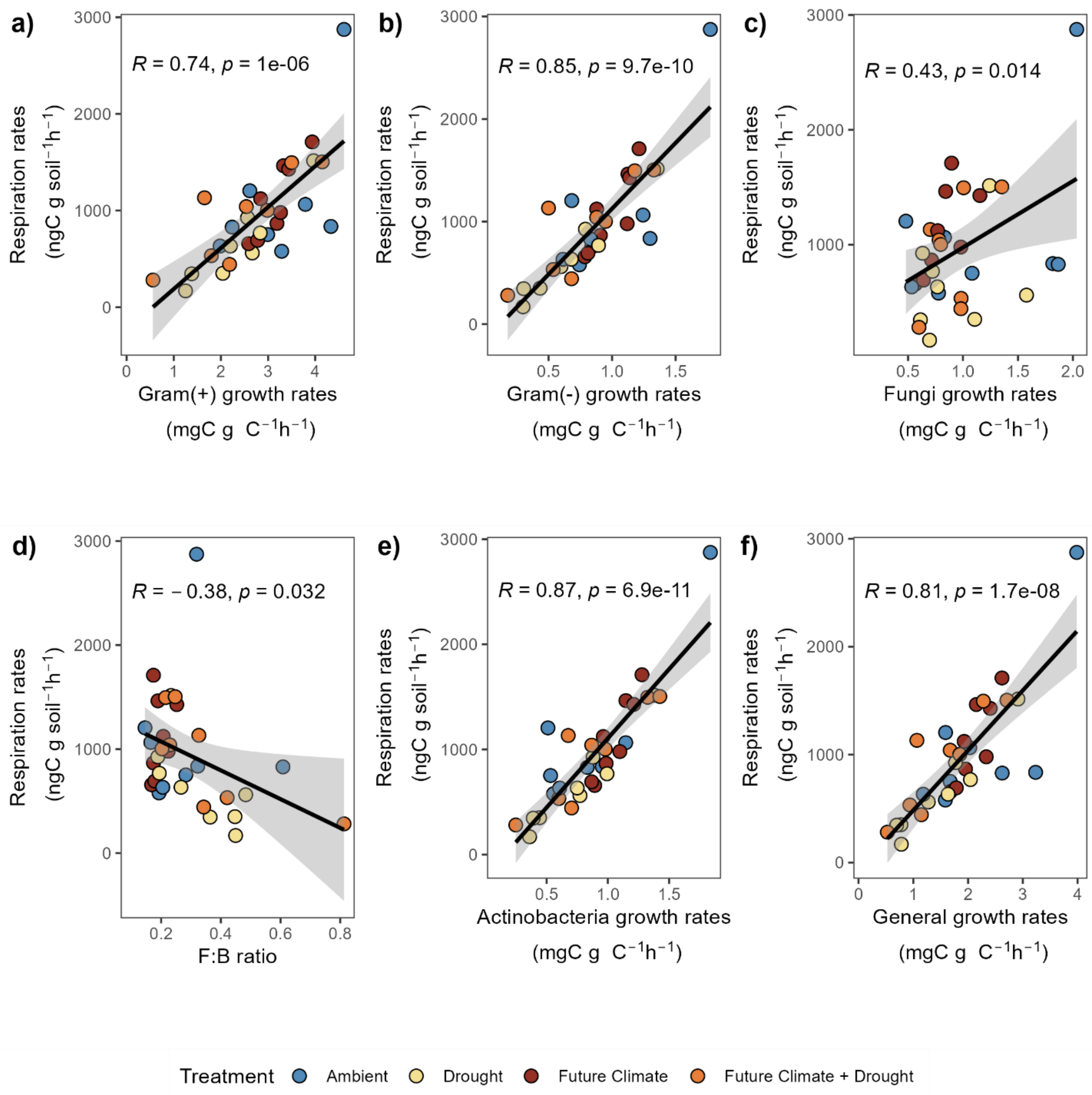


**Fig. S4.**

**Correlations between mass specific growth rates and total respiration rates** of gram positive (a), gram negative (b), fungi (c), actinobacteria (e) and general biomarkers (f). Correlation with funi to bacteria ratio is shown in panel d). Correlation were taken across all the time points mesured (Drought and Recovery). Pearson correlation coefficient (R) and p-value are reported on each graph. For all the graphs color indicates treatment (Ambient = blue, Drought = yellow, Future Climate = red, Future Climate + Drought = orange).

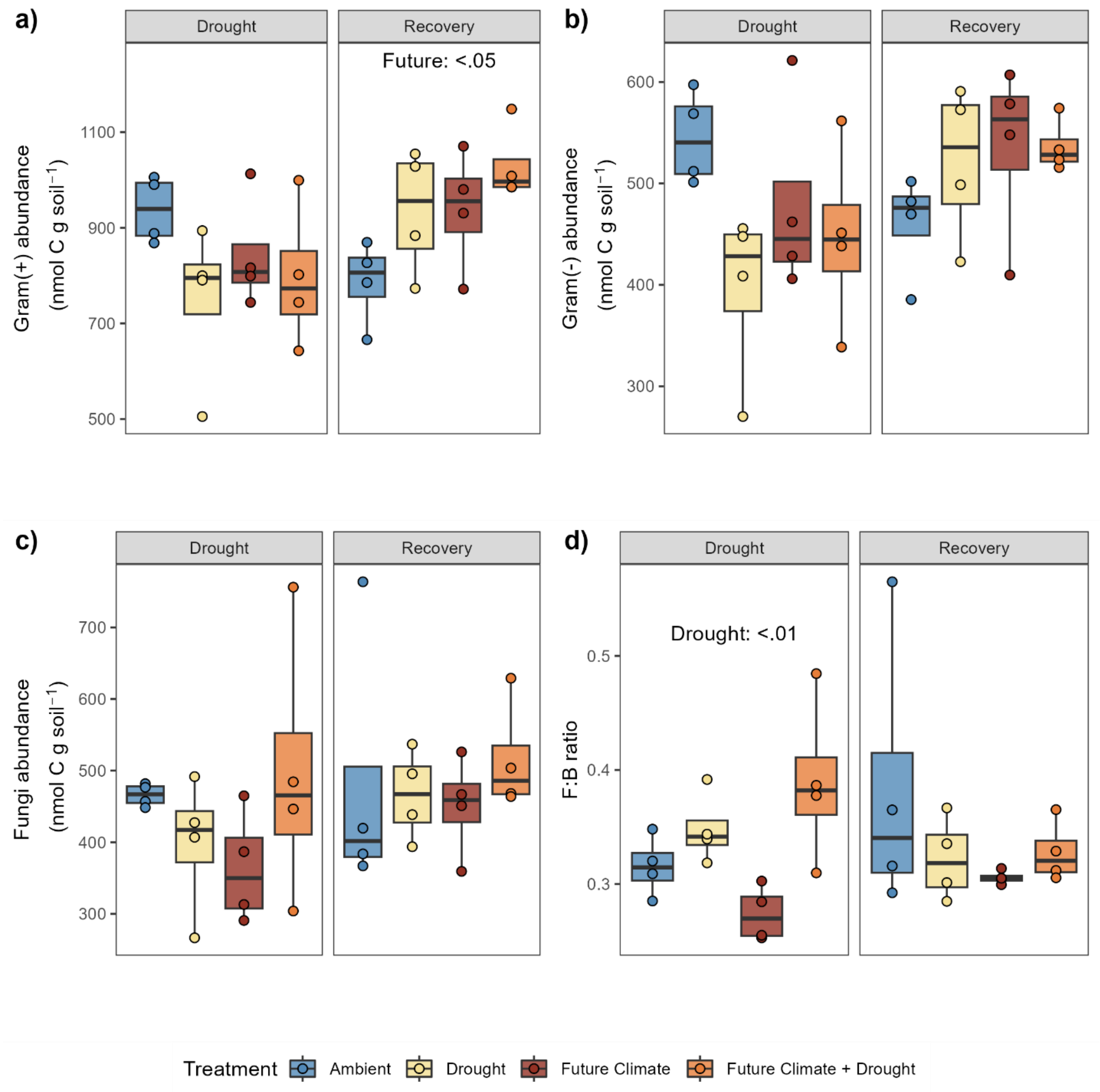


**Fig. S5.**

**PLFA-based biomass of microbial groups** for a) Gram positive b) Gram negative and c) fungi, divided by the time points measured (Drought and Recovery). Fungi to bacteria ratio is reported in d). Statistical results are reported only for significant p-values. Box centre line represents median, box limits the upper and lower quartiles, whiskers the 1.5x interquartile range, while separated points represent potential outliers. The sample size 'n' represents biologically independent samples (n=4). For all the graphs colour indicates treatment (Ambient = dark blue, Drought = light blue, Future Climate =red, Future Climate + Drought = orange).

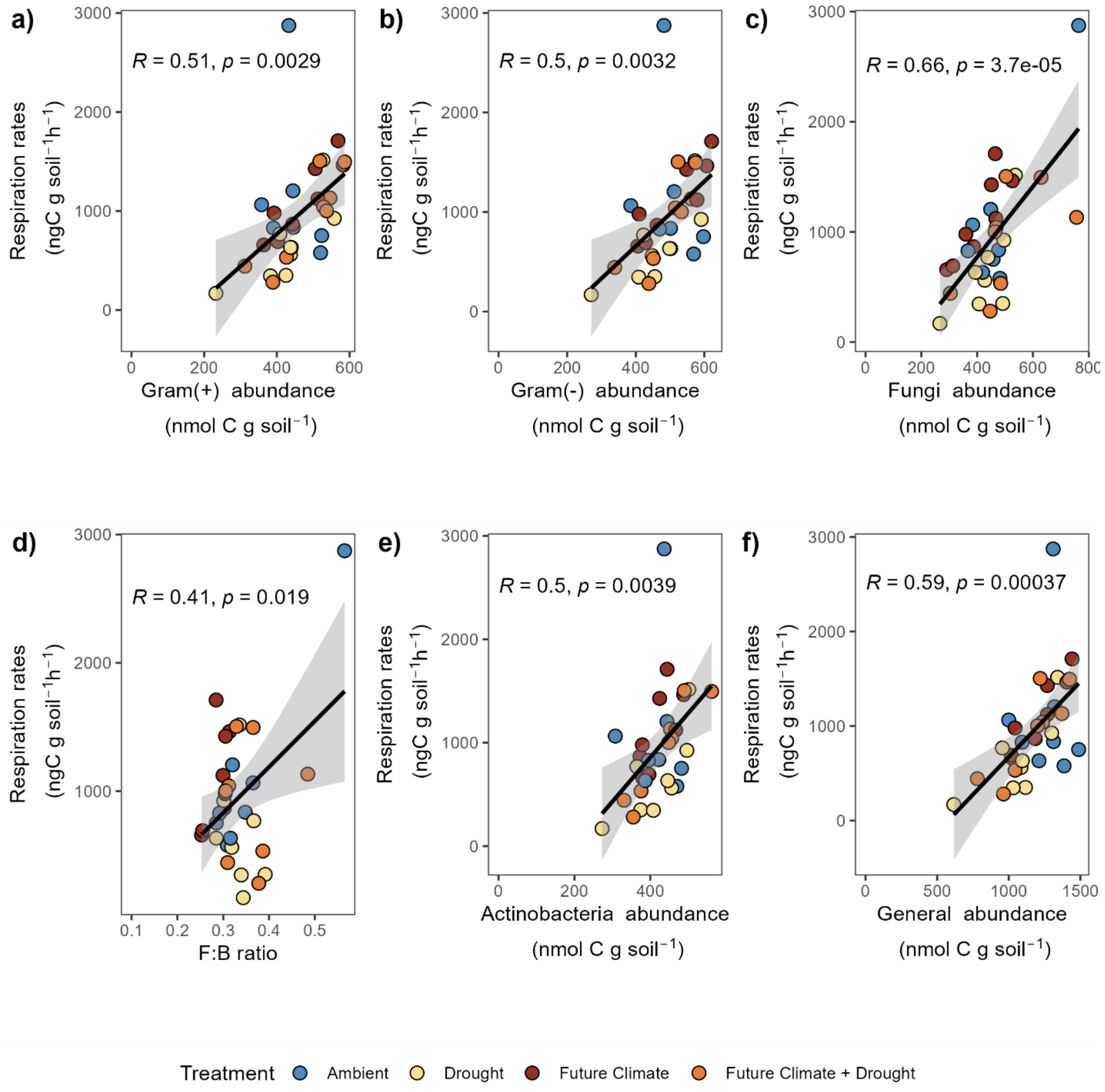


**Fig. S6.**

**Correlations between PLFA based microbial biomass separated by groups and total respiration** rates of a) Gram positive b) Gram negative c) fungi e) actinobacteria, f) and general biomarkers. Correlation with funi to bacteria ratio is shown in panel d). Correlation were taken across all the time points mesured (Drought and Recovery). Pearson correlation coefficient (R) and p-value are reported on each graph. For all the graphs color indicates treatment (Ambient = blue, Drought = yellow, Future Climate = red, Future Climate + Drought = orange).

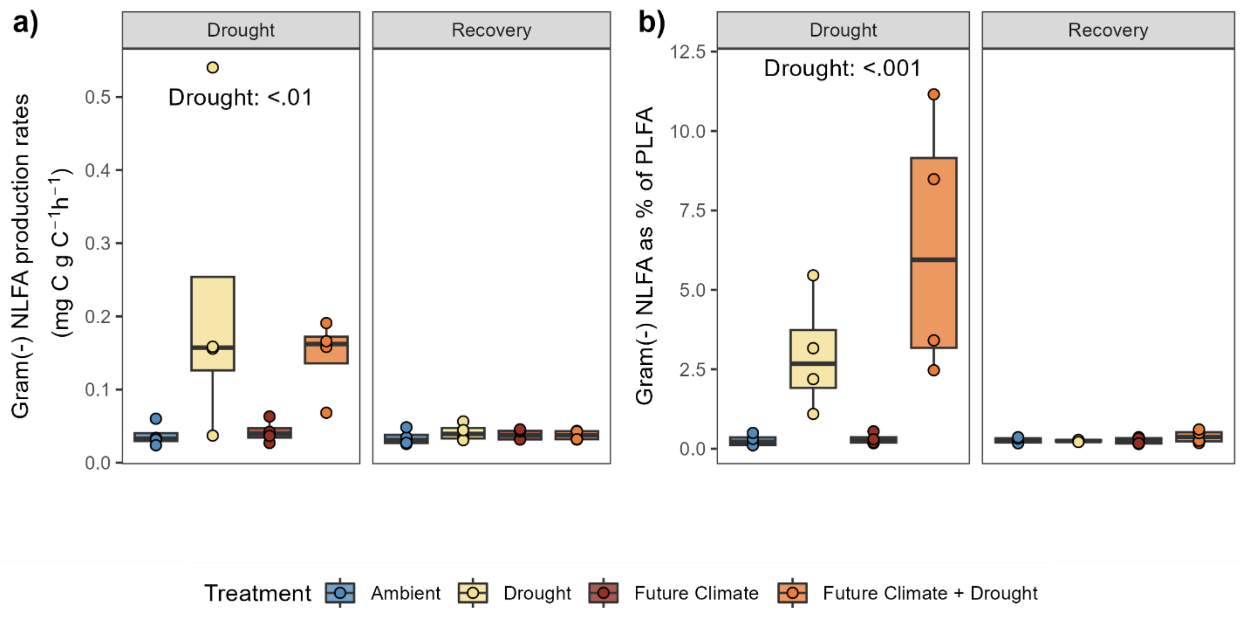


**Fig. S7.**

**Effects of drought and future climate on Gram negative NLFA biomarkers**. a) Production of gram negative specific NLFA biomarker and b) Investment in NLFA relative to the same Gram-negative PLFA specific biomarkers (shown as percent), divided by the time points measured (Drought and Recovery). Statistical results are reported for significant p-values (for a full report see Table S4). Box center line represents median, box limits the upper and lower quartiles, whiskers the 1.5x interquartile range, while separated points represent potential outliers. The sample size 'n' represents biologically independent samples (n=4). Color indicates treatment (Ambient = blue, Drought = yellow, Future Climate = red, Future Climate + Drought = orange).

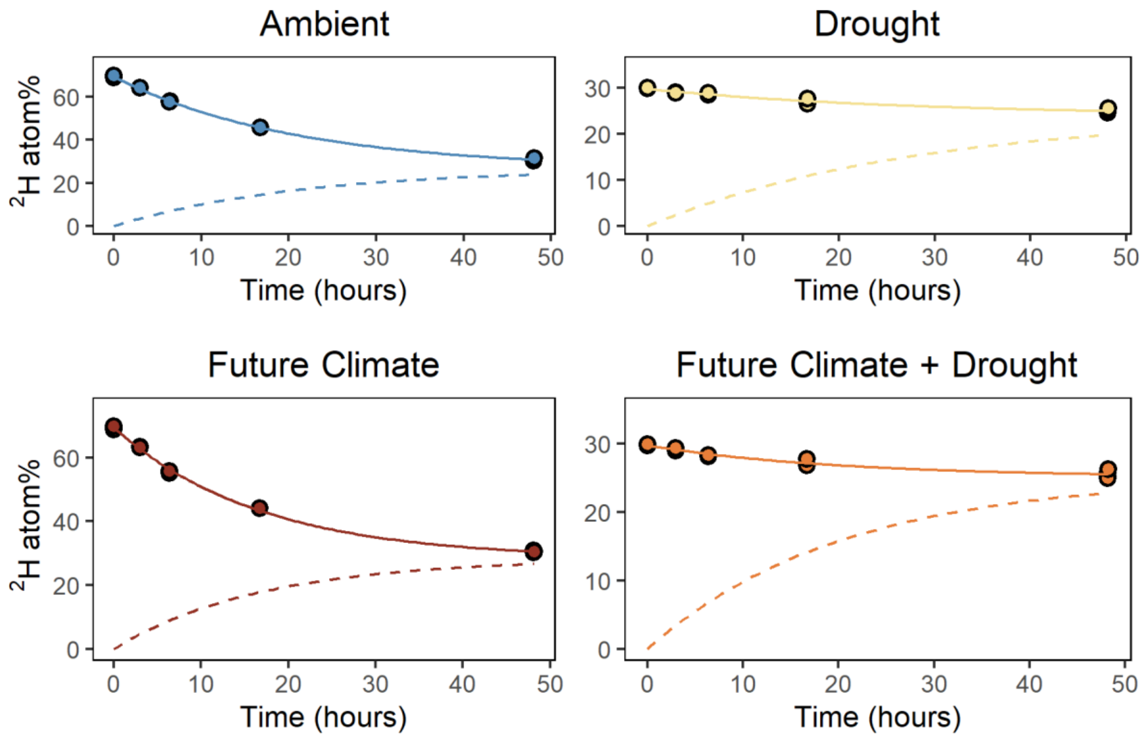


**Fig. S8.**

**Temporal dynamic of $^2$H equilibration with soil water.** Points represent measured $^2$H atom% values in the $^2$H labelled source water reaching isotopic equilibration rates of the external $^2$H labelled source water with soil water of the four treatments. Continuous lines represent the best fit model as used in Canarini et al. (2020) and described in Supplementary Materials and Methods. Dashed lines represent the model prediction of $^2$H kinetics in the soil water, generated from the data points of the labelled source water, as described in Canarini et al. (2020).

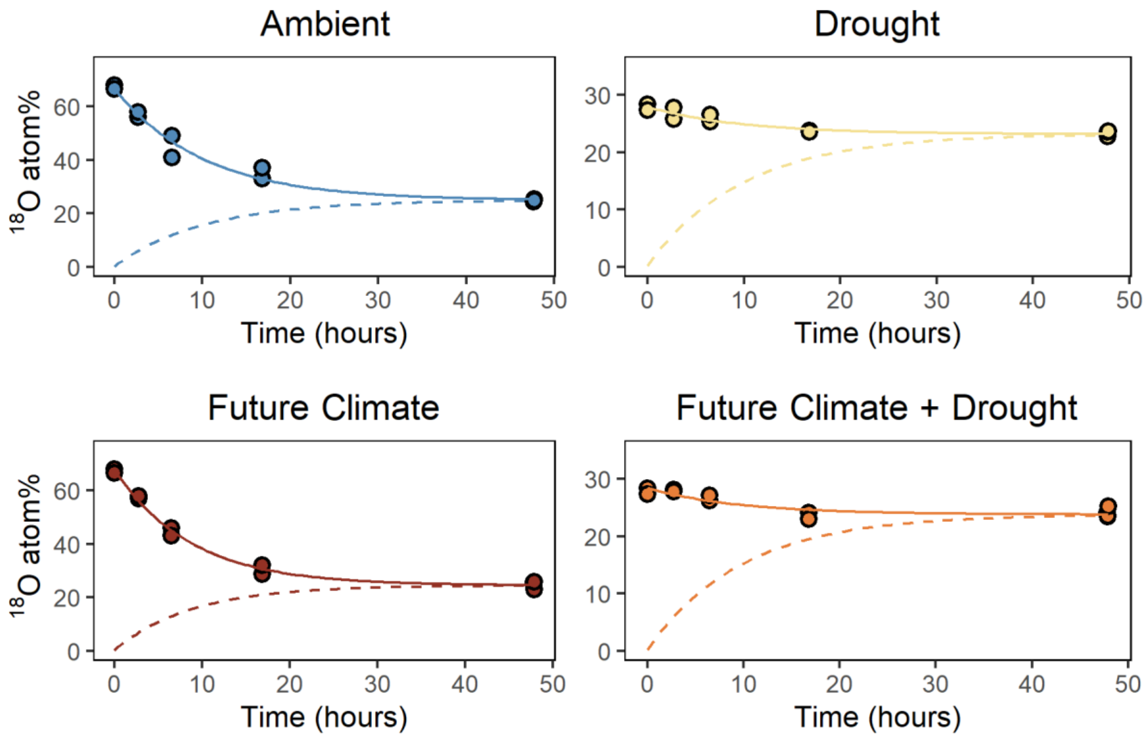


**Fig. S9.**

**Temporal dynamic of $^{18}O$ equilibration with soil water.** Points represent measured $^{18}O$ atom% values in the $^{18}O$ labelled source water reaching isotopic equilibration rates of the external $^{18}O$ labelled source water with soil water of the four treatments. Continuous lines represent the best fit model as used in Canarini et al. (2020) and described in Supplementary Materials and Methods. Dashed lines represent the model prediction of $^{18}O$ kinetics in the soil water, generated from the data points of the labelled source water, as described in Canarini et al. (2020).

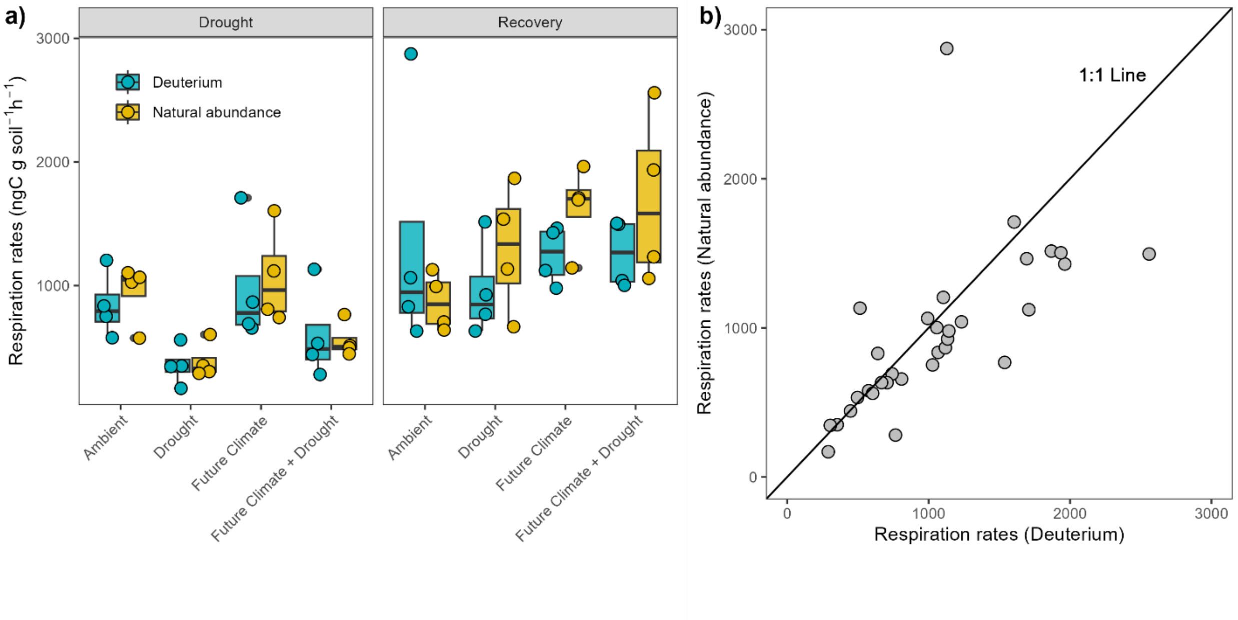


**Fig. S10.**

**Comparison of respiration rates determined samples incubated with non-labelled (natural abundance) vs. $^{2}H$ labelled water.** a) Values obtained for natural abundance samples (yellow) and labelled samples (blue) for all different treatments and time periods. Statistical analysis (paired t-test) revealed no significant differences. Box center line represents median, box limits the upper and lower quartiles, whiskers the 1.5x interquartile range, while separated points represent potential outliers. The sample size ‘n’ represents biologically independent samples (n=4). b) correlation between natural abundance samples and labelled samples around the 1:1 line.

**Table S1.**

**Effect of drought and future climate on mass-specific growth, mass-specific respiration and CUE determined by $^{2}H$- and $^{18}O$-vapor SIP, respectively.** We used linear mixed models to test effects of drought and the future climate change treatment and their interaction during 'Drought' and 'Recovery' on mass specific growth (mg C g microbial $C^{-1}$ $h^{-1}$) and microbial CUE; *p*-values < 0.05 are given in bold (*n* = 4 per treatment).

| | Growth | | | | | | | |
|---|---|---|---|---|---|---|---|---|
| | *'Drought'* | | | | *'Recovery'* | | | |
| | $^{2}H$-PLFA | | $^{18}O$-DNA | | $^{2}H$-PLFA | | $^{18}O$-DNA | |
| *Factor* | *F* | *p* | *F* | *p* | *F* | *p* | *F* | *p* |
| Drought | 15.061 | **0.002** | 48.878 | **<.0001** | 0.538 | 0.477 | 0.088 | 0.771 |
| Future Climate | 0.085 | 0.775 | 0.046 | 0.832 | 0.179 | 0.679 | 3.988 | 0.069 |
| Interaction | 0.004 | 0.945 | 0.117 | 0.737 | 0.319 | 0.582 | 0.0002 | 0.988 |

| | Respiration | | | | | | | |
|---|---|---|---|---|---|---|---|---|
| | *'Drought'* | | | | *'Recovery'* | | | |
| | $^{2}H$-PLFA | | $^{18}O$-DNA | | $^{2}H$-PLFA | | $^{18}O$-DNA | |
| Drought | 7.727 | **0.016** | 12.436 | **0.004** | 0.019 | 0.892 | 1.324 | 0.272 |
| Future Climate | 3.770 | 0.076 | 1.204 | 0.293 | 20.634 | **0.0007** | 27.394 | **0.0002** |
| Interaction | 0.531 | 0.479 | 0.001 | 0.970 | 1.508 | 0.242 | 0.079 | 0.782 |

| | CUE | | | | | | | |
|---|---|---|---|---|---|---|---|---|
| | *'Drought'* | | | | *'Recovery'* | | | |
| | $^{2}H$-PLFA | | $^{18}O$-DNA | | $^{2}H$-PLFA | | $^{18}O$-DNA | |
| *Factor* | *F* | *p* | *F* | *p* | *F* | *p* | *F* | *p* |
| Drought | 0.003 | 0.951 | 0.256 | 0.621 | 0.049 | 0.826 | 3.027 | 0.107 |
| Future Climate | 3.610 | 0.081 | 3.395 | 0.090 | 99.734 | **<0.0001** | 32.931 | **0.0001** |
| Interaction | 2.287 | 0.156 | 1.034 | 0.329 | 1.778 | 0.106 | 0.734 | 0.408 |

**Table S2.**

**Effects of drought and future climate on PLFA community composition (relative abundance) and relative PLFA-based growth (growth rates),** using a permanova based on Euclidean distance matrices as described in the results section for ‘Drought’ and ‘Recovery’. *p*-values < 0.05 are given in bold (*n* = 4 per treatment; Df = degrees of freedom).

| | *‘Drought’* | | | | | |
|---|---|---|---|---|---|---|
| | Abundance | | | Growth rates | | |
| | Df | $R^2$ | *p* | Df | $R^2$ | *p* |
| Drought | 1 | 0.297 | **0.001** | 1 | 0.405 | **0.001** |
| Future Climate | 1 | 0.139 | **0.019** | 1 | 0.246 | **0.001** |
| Interaction | 1 | 0.047 | 0.341 | 1 | 0.065 | **0.040** |
| | *‘Recovery’* | | | | | |
| | Abundance | | | Growth rates | | |
| | Df | $R^2$ | *p* | Df | $R^2$ | *p* |
| Drought | 1 | 0.181 | **0.002** | 1 | 0.140 | **0.034** |
| Future Climate | 1 | 0.119 | **0.012** | 1 | 0.070 | 0.330 |
| Interaction | 1 | 0.094 | 0.064 | 1 | 0.077 | 0.281 |

**Table S3.**

**Statistical results of individual group mass-specific growth rates obtained by $^2H$ incorporation into PLFA.** Statistical results of drought, climate change treatment (Future Climate) and their interaction during 'Drought' and 'Recovery' for individual group mass specific growth rates (mg C g microbial $C^{-1}$ $h^{-1}$). Values are derived from a linear mixed effect model (as described in materials and method) for each sampling date and separately for Fungi, Gram negative and Gram positive. *p* values < 0.05 are given in bold (*n* = 4 in each treatment).

| | *'Drought'* | | | | | |
|---|---|---|---|---|---|---|
| *Factor* | Drought | | Future Climate | | Interaction | |
| | *F* | *p* | *F* | *p* | *F* | *p* |
| Fungi | 0.176 | 0.683 | 1.426 | 0.256 | 0.327 | 0.578 |
| Gram negative | 23.830 | **<0.0001** | 0.414 | 0.532 | 0.015 | 0.905 |
| Gram positive | 20.468 | **0.0007** | 0.486 | 0.498 | 0.023 | 0.881 |
| F:B ratio | 31.299 | **<0.0001** | 0.712 | 0.415 | 0.961 | 0.346 |
| Actinobacteria | 7.232 | **0.019** | 2.756 | 0.122 | 1.220 | 0.290 |
| General | 29.215 | **0.0002** | 0.038 | 0.846 | 0.000 | 0.999 |

| | *'Recovery'* | | | | | |
|---|---|---|---|---|---|---|
| *Factor* | Drought | | Future Climate | | Interaction | |
| | *F* | *p* | *F* | *p* | *F* | *p* |
| Fungi | 1.075 | 0.320 | 0.319 | 0.583 | 1.615 | 0.228 |
| Gram negative | 0.194 | 0.668 | 0.038 | 0.849 | 0.181 | 0.678 |
| Gram positive | 0.052 | 0.823 | 0.027 | 0.871 | 0.174 | 0.683 |
| F:B ratio | 0.597 | 0.454 | 0.589 | 0.457 | 0.929 | 0.345 |
| Actinobacteria | 0.034 | 0.856 | 0.006 | 0.936 | 0.207 | 0.656 |
| General | 0.390 | 0.543 | 0.270 | 0.612 | 0.171 | 0.686 |

**Table S4.**

**Impacts of drought and future climate on NLFA production.** Statistical results of drought, climate change treatment (Future Climate) and their interaction during 'Drought' and 'Recovery' for NLFA production (ngC g soil$^{-1}$ h$^{-1}$) tested by linear mixed effect model (as described in materials and method) for each sampling date separately for fungi, gram negative and total (sum of all) biomarkers. *p* values < 0.05 are given in bold (*n* = 4 per treatment).

| | Fungi (ng C g C$^{-1}$ h$^{-1}$) | | Gram negative (ng C g C$^{-1}$ h$^{-1}$) | | Fungi (% of PLFA) | | Gram negative (% of PLFA) | |
|---|---|---|---|---|---|---|---|---|
| ***'Drought'*** | | | | | | | | |
| *Factor* | *F* | *p* | *F* | *p* | *F* | *p* | *F* | *p* |
| Drought | 22.485 | **0.0005** | 16.717 | **0.015** | 87.339 | **<.0001** | 66.491 | **<.0001** |
| Future Climate | 0.373 | 0.552 | 0.004 | 0.948 | 0.527 | 0.481 | 2.249 | 0.159 |
| Interaction | 0.147 | 0.707 | 0.121 | 0.733 | 0.200 | 0.662 | 0.511 | 0.488 |
| ***'Recovery'*** | | | | | | | | |
| | Fungi (ng C g C$^{-1}$ h$^{-1}$) | | Gram negative (ng C g C$^{-1}$ h$^{-1}$) | | Fungi (% of PLFA) | | Gram negative (% of PLFA) | |
| *Factor* | *F* | *p* | *F* | *p* | *F* | *p* | *F* | *p* |
| Drought | 0.014 | 0.905 | 0.580 | 0.460 | 0.607 | 0.450 | 0.635 | 0.442 |
| Future Climate | 0.408 | 0.534 | 0.0001 | 0.989 | 1.215 | 0.291 | 0.396 | 0.540 |
| Interaction | 0.005 | 0.940 | 0.760 | 0.400 | 0.300 | 0.593 | 1.088 | 0.317 |
